# Demonstration of the potential of environmental DNA as a tool for the detection of avian species

**DOI:** 10.1101/199208

**Authors:** Masayuki Ushio, Koichi Murata, Tetsuya Sado, Isao Nishiumi, Masamichi Takeshita, Wataru Iwasaki, Masaki Miya

## Abstract

Birds play unique functional roles in the maintenance of ecosystems, such as pollination and seed dispersal, and thus monitoring bird communities (e.g., monitoring bird species diversity) is a first step towards avoiding undesirable consequences of anthropogenic impacts on bird communities. In the present study, we hypothesized that birds, regardless of their main habitats, must have frequent contact with water and that tissues that contain their DNA that persists in the environment (environmental DNA; eDNA) could be used to detect the presence of avian species. To this end, we applied a set of universal PCR primers (MiBird, a modified version of fish/mammal universal primers) for metabarcoding avian eDNA. We confirmed the versatility of MiBird primers by performing *in silico* analyses and by amplifying DNAs extracted from bird tissues. Analyses of water samples from zoo cages of birds with known species composition suggested that the use of MiBird primers combined with the Illumina MiSeq platform could successfully detect avian species from water samples. Additionally, analysis of water samples collected from a natural pond detected five avian species common to the sampling areas. The present findings suggest that avian eDNA metabarcoding would be a complementary detection/identification tool in cases where visual detection and identification of bird species is difficult.

## I. INTRODUCTION

Environmental DNA (eDNA) is genetic material that persists in an environment and is derived from organisms living there, and researchers have recently been using eDNA to detect the presence of macro-organisms, including those living in aquatic/semiaquatic ecosystems [1–5]. For example, several fish species inhabiting a river can be detected by amplifying and sequencing DNA fragments extracted from water samples [6] by using methodologies such as quantitative PCR and eDNA metabarcoding. Quantitative PCR requires the design of species-specific PCR primers and enables quantitative measurements of eDNA of target species [3, 4, 7, 8] while eDNA metabarcoding, which has been becoming a common methodology in eDNA studies, uses a universal primer set and high-throughput sequencer (e.g., Illumina MiSeq) to enable qualitative detection of eDNA of multiple species belonging to a target taxon [1, 2, 9–11] (but see [12]).

Although earlier studies mainly focused on detecting fish/amphibian species (i.e., organisms that have close associations with water), recent studies have shown that eDNA can be used to detect a diverse group of animals, including mammals [9, 13], reptiles [14] and arthropods [10]. Detecting the presence of animals is possible even if their habitats are terrestrial [9, 10, 13] because animals must have, in general, frequent opportunities to contact water in order to live. The findings of these recent studies imply that any organism, regardless of its main habitat, can potentially be detected by using eDNA if we can design suitable primers that enable amplification and identification of DNA fragments of target organisms and if we can collect appropriate media that contain eDNA.

Wild birds represent an important part of the biodiversity in ecosystems, and they play a unique role in the maintenance of ecosystem functions. For example, in forest ecosystems, birds can contribute to maintenance of the tree community by seed dispersal and pollination, and to the reduction of herbivory by predation upon insect herbivores [15–17]. However, recent increases in anthropogenic impacts on ecosystems, e.g., urbanization and habitat fragmentation, drive substantial declines in bird species diversity [18, 19], which could have impacts on the ecological functions of birds. Monitoring bird species diversity is required for detecting such declines, and such detection is necessary for avoiding undesirable consequences in ecosystem functions due to the loss of biodiversity. To monitor bird species diversity, visual census is one of the most common methods [20], and considering the higher visibility of birds than that of fish and forest mammals, visual census is generally a successful method. However, if an alternative method can overcome limitations of visual census, such as low visibility at night or in a dense forest, and eliminate the requirement for taxonomic identification skill under field conditions, that method could be complementarily used for monitoring bird species diversity.

In the present study, we tested the potential of eDNA as a tool for the detection of avian species. To this end, we modified a previously reported universal primer set for fish/mammals, (MiFish/MiMammal [1, 9]) such that the primer set accommodated bird-specific variations, and conducted avian eDNA metabarcoding. During the primer design, we did not try to eliminate the capability of detecting mammalian species, because simultaneous detection of mammals along with birds may be advantageous, especially for ecologists who are interested in co-occurrence patterns and potential interactions among various animal species. We performed a series of analyses to test the versatility of the designed primers: *In silico* examinations of the primers, amplification of extracted tissue DNAs of birds belonging to various taxa, and field tests by analyzing water samples from zoo cages containing birds of known species composition. Additionally, we briefly examined the usefulness of the new primer set using water samples from field samples with unknown bird species composition.

## II. METHODS

All of the critical information of our study is described below, but is also listed in Table S1 to facilitate comparisons with other studies, following the recommendations of Goldberg et al. [21].

### A. Primer design

To facilitate design based on comparisons of diverse avian sequences, we first batch downloaded 410 avian sequences from RefSeq (https://www.ncbi.nlm.nih.gov/refseq/) on June 9, 2015. Then, a base composition for a selected position in the conservative region was shown in Mesquite [22]. The base compositions in selected characters were manually recorded in a spreadsheet for the primer design. In the primer design process, we considered a number of technical tips that enhance the primer annealing to the template without the use of degenerate bases [23]: primers include some G/C at the 3’-ends to strengthen primer-template annealing at this position, but a string of either Gs or Cs at the 3’-end should be avoided: considering the unconventional base pairing in the T/G bond, the designed primers use G rather than A when the template is variably C or T, and T rather than C when the template is A or G; G*/*C contents of the primers fall between 40 and 60%, with an almost identical melting temperature (Tm). Tm was calculated using a nearest-neighbour thermodynamic model implemented in OLIGOCALC [24].

We designed our primers by modifying previously developed MiFish/MiMammal primers [1, 9], which corresponded to regions in the mitochondrial 12SrRNA gene (insert length = *ca*. 171 bp), and we named our primers MiBird-U (“U” indicates “universal”). Primer sequences with MiSeq adaptors (for the first- and second-round PCR) are listed in Table 1.

**Table 1:**
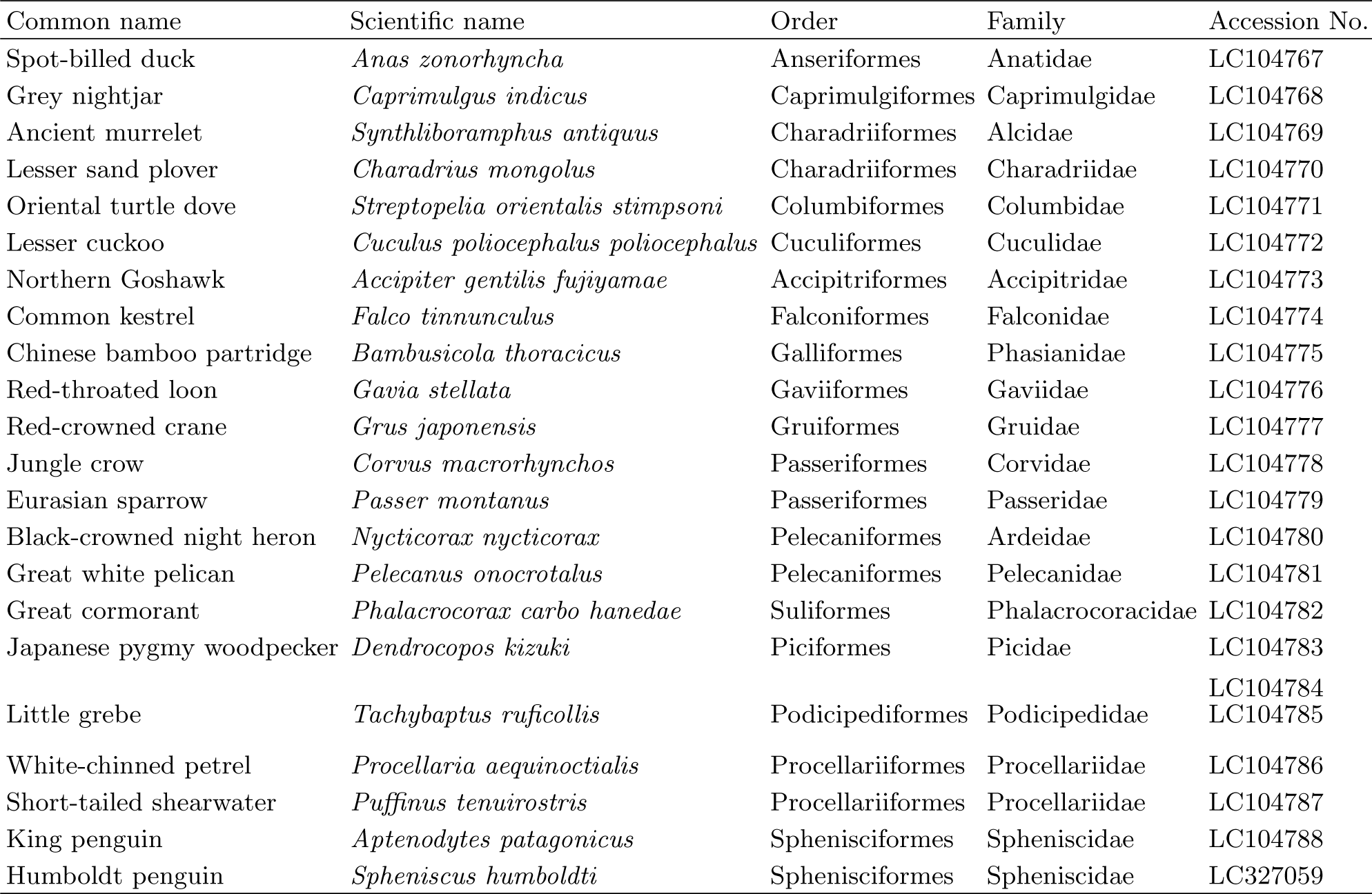
Detailed information for MiBird primers

### B. **In silico** evaluation of interspecific variation of MiBird sequences

The binding capacity of MiBird-U primers was computationally evaluated using the batch-downloaded 410 avian sequences. Using custom Ruby and Python scripts, the number of mismatches between MiBird-U primers and the 410 avian sequences was calculated.

Interspecific differences within the amplified DNA sequences are required for assignment of taxonomic categories. Levels of interspecific variation in the target region (hereafter called ’MiBird sequence’) across different taxonomic groups of birds were computationally evaluated using the 410 downloaded avian sequences. Among the sequences of the 410 avian, species with the deletion of primer regions (*Hemignathus munroi*, *Loxops coccineus* and *Arborophila rufipectus*) were excluded, and 407 MiBird sequences were extracted and subjected to calcula-tion of pairwise edit distances using custom Python scripts. Pairwise inter-species edit distances were calculated for all species pairs, and pairwise intergenus edit distances were calculated for pairs of species belonging to different genera. The edit distance quantifies dissimilarity of sequences in bioinformatics and is defined as the minimum number of single-nucleotide substitutions, insertions or deletions that are required to transform one sequence into the other.

In addition, the binding capacity and the levels of interspecific variations of the target region were further evaluated using ’primerTree’ package [25] of R version 3.3.1 [26]. Briefly, primerTree performs the following analysis: (1) *In silico* PCR against sequences in the NCBI database; (2) retrieval of DNA sequences predicted to be amplified; (3) taxonomic identification of these sequences; (4) multiple DNA sequence alignment; (5) reconstruction of a phylogenetic tree and (6) visualization of the tree with taxonomic annotation. Thus, by using primerTree package, species whose sequences can be amplified, phylogenetic relationships among these amplified species, and interspecific variations in the amplified sequences are rapidly visualized. Further information and instructions for the primerTree package can be found in Cannon et al. [25].

### C. Primer testing with extracted DNA

We tested the versatility of MiBird-U (no adapter sequences) using DNA extracted from 22 species representing major groups of birds (Table 2). Double-stranded DNA concentrations from those samples were measured with a NanoDrop Lite spectrophotometer (Thermo Fisher Scientific, Wilmington, DE, USA) and the extracted DNA was diluted to 15 ng *μ*l ^-1^ using Milli-Q water. PCR was carried out with 30 cycles of a 15 *μ*l reaction volume containing 4.5 *μ*l sterile distilled H_2_O, 7.5 *μ*l 2 × Gflex PCR Buffer (Mg^2^+, dNTPs plus) (Takara, Otsu, Japan), 0.7 *μ*l of each primer (5 *μ*M), 0.3 *μ*l Taq polymerase (Tks Gflex DNA Polymerase; Takara) and 1.2 *μ*l template. The thermal cycle profile after an initial 1 min denaturation at 94°C was as follows: denaturation at 98°C for 10 s; annealing at 50°C for 10 s; and extension at 68°C for 10 s with a final extension at the same temperature for 7 min.

**Table 2:**
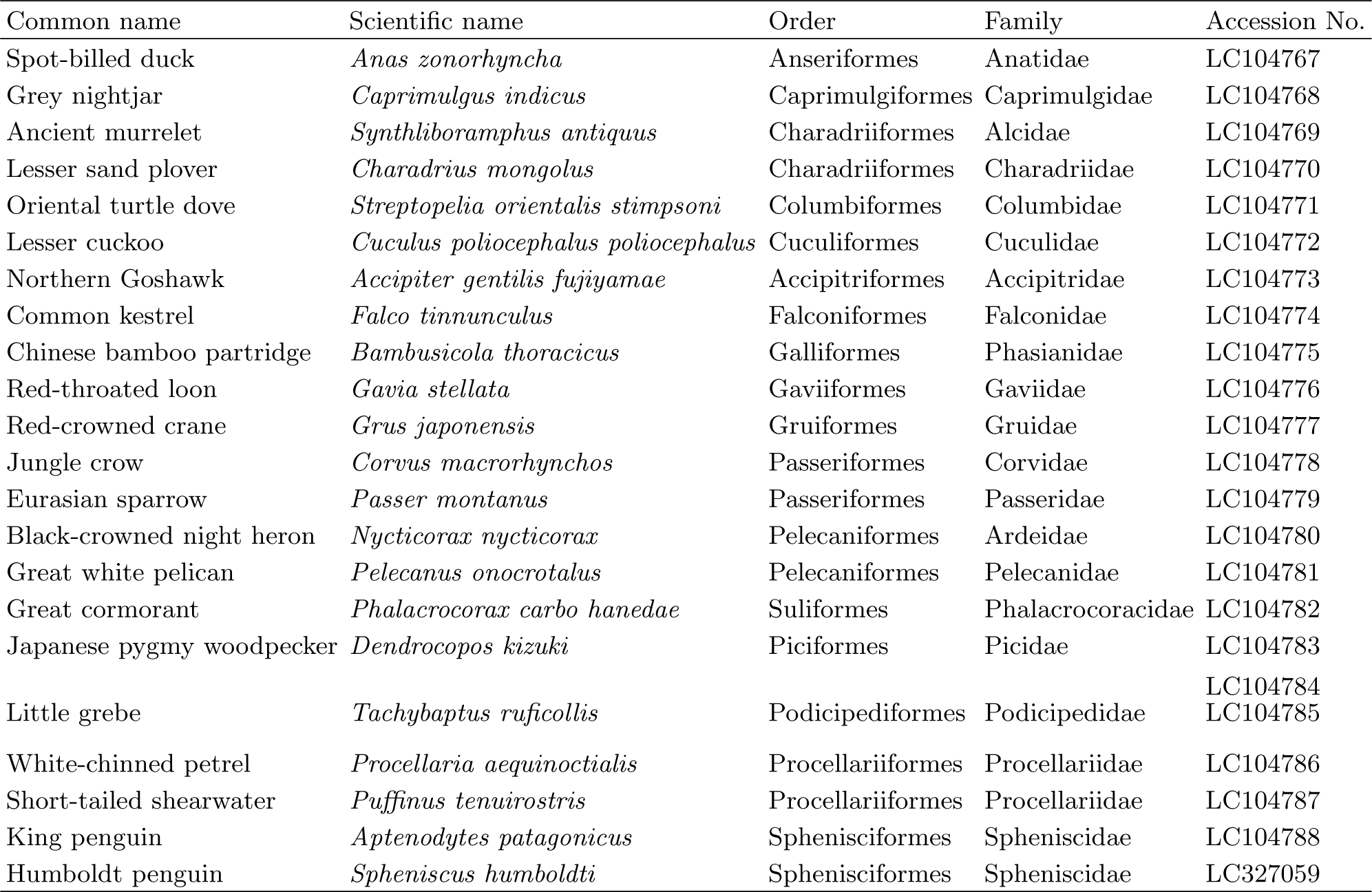
Extract DNAs used to test the performance of the developed primer set

### D. Study site and water sampling for primer testing with eDNA from zoo samples

To test the versatility of the newly designed primers for metabarcoding avian eDNA, we sampled water from cages on 13 December 2016 in Yokohama Zoological Gardens ZOORASIA, Yokohama, Japan (35°29’42” N, 139°31’35” E), where we previously tested the usefulness of a universal primer set targeting mammals [9]. We chose the zoo as a sampling site because the information about avian species in a cage is precisely known, and because the zoo rears diverse taxonomic groups of animals (i.e., > 100 animal species, including many mammals and birds). Thirteen cages, in which diverse taxonomic groups of birds were reared, were selected as sampling places (Table 3). Most of the target species were kept separately, but ruddy shelduck (*Tadorna ferruginea*) were kept in a walk through bird cage (hereafter,“the bird cage”) with other bird species (i.e., Lady Amherst’s pheasant [*Chrysolophus amherstiae*], Temminck’s tragopan [*Tragopan temminckii*], Victoria crowned pigeon [*Goura victoria*] and mandarin duck [*Aix galericulata*]). Note that different individuals of Lady Amherst’s pheasant, Temminck’s tragopan, Victoria crowned pigeon and mandarin duck were separately kept in different cages from the bird cage, and that a water sample of each bird species was collected from each cage.

**Table 3:** Classification of the target bird species in the Zoorasia experiment

| Common name | Species name | Order | Family |
| --- | --- | --- | --- |
| Steller’s sea eagle | *Haliaeetus pelagicus* | Accipitriformes | Accipitridae |
| Black-tailed gull | *Larus crassirostris* | Charadriiformes | Laridae |
| Capercaillie | *Tetrao urogallus* | Galliformes | Tetraonidae |
| Lady Amherst’s pheasant | *Chrysolophus amherstiae* | Galliformes | Phasianidae |
| Ruddy shelduck | *Tadorna ferruginea* | Anseriformes | Anatidae |
| Temminck’s tragopan | *Tragopan temminckii* | Galliformes | Phasianidae |
| Victoria crowned pigeon | *Goura victoria* | Columbiformes | Columbidae |
| Mandarin duck | *Aix galericulata* | Anseriformes | Anatidae |
| Humboldt penguin | *Spheniscus humboldti* | Sphenisciformes | Spheniscidae |
| Snowy owl | *Bubo scandiacus* | Strigiformes | Strigidae |
| Oriental white stork | *Ciconia boyciana* | Ciconiiformes | Ciconiidae |
| White-naped crane | *Grus vipio* | Gruiformes | Gruidae |
| Common crane | *Grus grus* | Gruiformes | Gruidae |
| Southern ground hornbill | *Bucorvus cafer* | Coraciiformes | Bucerotidae |
| Harris’s hawk | *Parabuteo unicinctus* | Accipitriformes | Accipitridae |
| Emu | *Dromaius novaehollandiae* | Struthioniformes | Casuariidae |

Each 100 200 ml water sample was collected through a sterile 0.45-*μ*m-pore Sterivex filter (Merck Millipore, Darmstadt, Germany) using a sterile 50- mL syringe (TERUMO, Tokyo, Japan). After the filtration, approximately 2 ml of RNAlater (Ther-moFisher Scientific, Waltham, Massachusetts, USA) was injected into the Sterivex cartridge, and the filtered water samples were stored at 4°C for up to one day until further processing. Three negative controls (distilled water) were taken to the zoo to monitor contaminations during water sampling, filtration and transport.

In addition to the survey in the zoo, we collected water samples from a pond adjacent to the Natural History Museum and Institute, Chiba (35°35’59” N, 140°8’18” E) to test the potential effectiveness of the MiBird primers under a field condition with unknown bird species composition. Water collections at the pond were performed in the same way as those performed in the zoo.

### E. DNA extraction

The Sterivex filter cartridges were taken back to the laboratory, and DNA was extracted from the filters using a DNeasy Blood and Tissue Kit (Qiagen, Hilden, Germany) following a protocol described and illustrated in Miya et al. [27]. Briefly, the RNAlater-supplemented solution was dried under a vacuum using the QIAvac system (Qiagen, Hilden, Germany). Proteinase-K solution (20 *μ*l), phosphate buffered saline (PBS) (220 *μ*l) and buffer AL (200 *μ*l) were mixed, and 440 *μ*l of the mixture was added to each filter cartridge. The materials on the filter cartridges were subjected to cell-lysis conditions by incubating the filters on a rotary shaker (at a speed of 20 rpm) at 50°C for 20 min. The incubated mixture was trans ferred into a new 2-ml tube, and the collected DNA was purified using a DNeasy Blood and Tissue Kit following the manufacturer’s protocol. After the purification, DNA was eluted using 100 *μ*l of the elution buffer provided with the kit.

### F. Paired-end library preparation

Prior to the library preparation, work-spaces and equipment were sterilized. Filtered pipet tips were used, and separation of preand post-PCR sam-ples was carried out to safeguard against cross-contamination. We also employed two negative controls (i.e., PCR negative controls) to monitor contamination during the experiments.

The first-round PCR (first PCR) was carried out with a 12-*μ*l reaction volume containing 6.0 *μ*l of 2 × KAPA HiFi HotStart ReadyMix (KAPA Biosystems, Wilmington, WA, USA), 0.7 *μ*l of MiBird primer (5*μ*M primer F*/*R, w*/* adaptor and six random bases; Table 1), 2.6 *μ*l of sterilized distilled H_2_O and 2.0 *μ*l of template. The thermal cycle profile after an initial 3 min denaturation at 95°C was as follows (35 cycles): denaturation at 98°C for 20 s; annealing at 65°C for 15 s; and extension at 72°C for 15 s, with a final extension at the same temperature for 5 min. We performed triplicate first-PCR, and these replicate products were pooled in order to mitigate the PCR dropouts. The pooled first PCR products were purified using AMPure XP (PCR product: AMPure XP beads = 1:0.8; Beckman Coulter, Brea, Califor-nia, USA). The pooled, purified, and 10-fold diluted first PCR products were used as templates for the second-round PCR.

The second-round PCR (second PCR) was carried out with a 24-*μ*l reaction volume containing 12 *μ*l of 2 × KAPA HiFi HotStart ReadyMix, 1.4 *μ*l of each primer (5 *μ*M primer F*/*R; Table 1), 7.2 *μ*l of sterilized distilled H_2_O and 2.0 *μ*l of template. Different combinations of forward and reverse indices were used for different templates (samples) for massively parallel sequencing with MiSeq. The thermal cycle profile after an initial 3 min denaturation at 95°C was as follows (12 cycles): denaturation at 98°C for 20 s; combined annealing and extension at 72°C (shuttle PCR) for 15 s, with a final extension at 72°C for 5 min.

The indexed second PCR products were mixed at equimolar concentrations to produce equivalent sequencing depth from all samples and the pooled library was purified using AMPure XP. Target-sized DNA of the purified library (*ca*. 370 bp) was excised using E-Gel SizeSelect (ThermoFisher Scientific, Waltham, MA, USA). The double-stranded DNA concentration of the library was quantified using a Qubit dsDNA HS assay kit and a Qubit fluorometer (ThermoFisher Scientific, Waltham, MA, USA). The double-stranded DNA concentration of the library was then adjusted to 4 nM using Milli-Q water and the DNA was applied to the MiSeq platform (Illumina, San Diego, CA, USA). The sequencing was performed using a MiSeq Reagent Kit Nano v2 for 2 × 150 bp PE (Illumina, San Diego, CA, USA). Note that although our MiSeq run yielded approximately 1,000,000 reads (an average number of reads for the MiSeq Reagent Kit Nano v2 for 2 × 150 bp PE), the total number of raw reads reported in this study is 633,375 and the rest of the reads correspond to those from other studies.

### G. Data processing and taxonomic assignment

The overall quality of the MiSeq reads was evaluated using the programs FASTQC (available from http://www.bioinformatics.babraham.ac.uk/) and SUGAR [28]. After confirming the lack of technical errors in the MiSeq sequencing, low-quality tails were trimmed from each read using DynamicTrim.pl from the SOLEXAQA software package [29] with a cut-off threshold set at a Phred score of 10 (= 10^-1^ error rate). The tail-trimmed pair-end reads were assembled using the software FLASH with a minimum overlap of 10 bp. The assembled reads were further filtered by custom Perl scripts in order to remove reads with either ambiguous sites or those showing unusual lengths compared to the expected size of the PCR amplicons. Finally, the software TagCleaner [30] was used to remove primer sequences with a maximum of three-base mismatches and to transform the FASTQ format into FASTA.

The pre-processed reads from the above custom pipeline were dereplicated using UCLUST [31], with the number of identical reads added to the header line of the FASTA formatted data file. Those sequences represented by at least 10 identical reads were subjected to the downstream analyses, and the remaining under-represented sequences (with less than 10 identical reads) were subjected to pairwise alignment using UCLUST. If the latter sequences observed for less than 10 reads showed at least 99% identity with one of the former reads (one or two nucleotide differences), they were operationally considered as identical (owing to sequencing or PCR errors and/or actual nucleotide variations in the populations).

The processed reads were subjected to local BLASTN searches [32] against a custom-made database. The custom database was generated by downloading all whole mitogenome sequences from Sarcopterygii deposited in NCBI Organelle Genome Resources (http://www.ncbi.nlm.nih.gov/genomes/OrganelleResource.cgifftaxid=8287). As of 15 March 2016, this database covered 1,881 species across a wide range of families and genera (including birds, mammals, reptiles and amphibians). In addition, the custom database was supplemented by all whole and partial fish mitogenome sequences deposited in MitoFish [33] in order to cover fish de-tection (note that MiBird primers amplify fish sequences as well; see Fig. 1).

**Figure 1:**
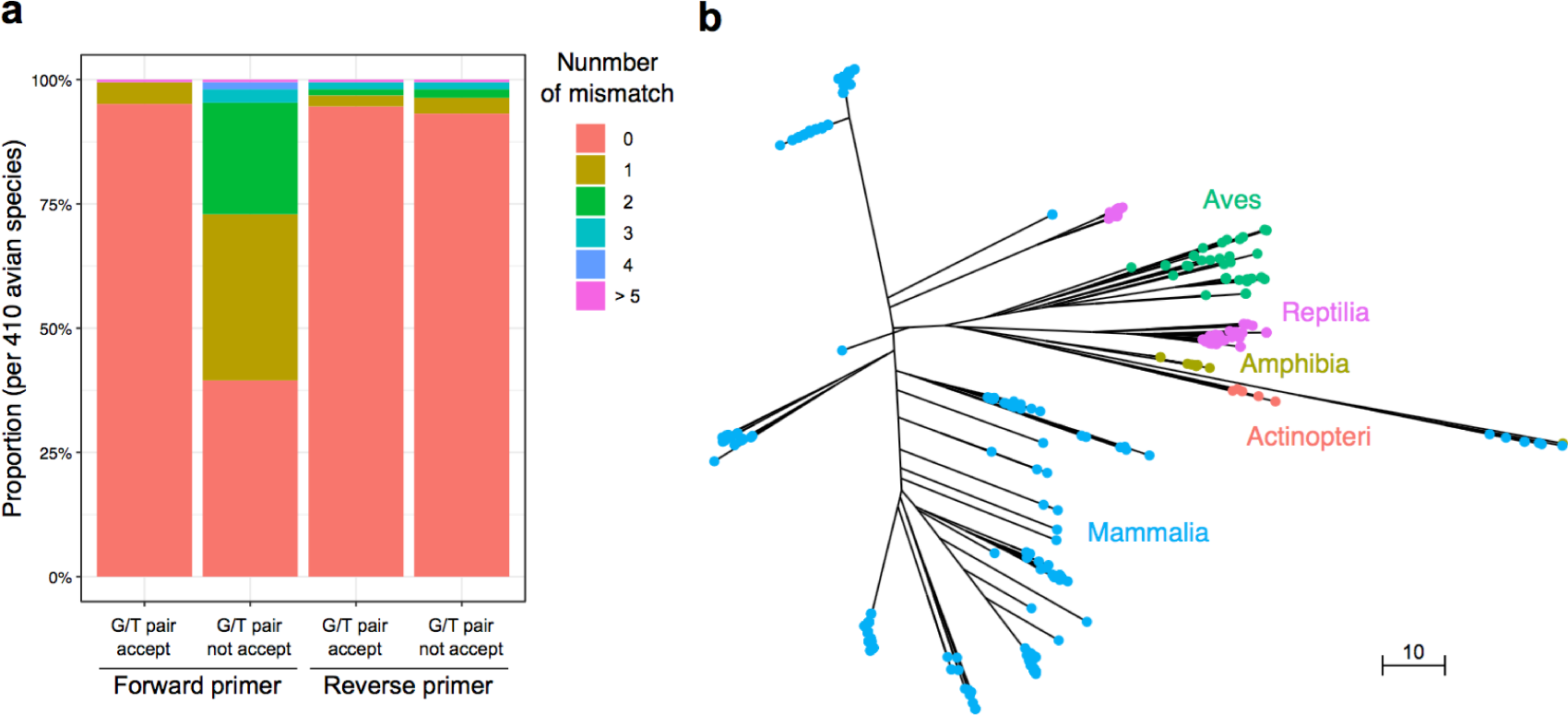
Results of *in silico* evaluations of MiBird-U primers. Binding capacity of MiBird-U primers (**a**). y-axis represents the proportion of avian species that showed 0, 1, 2, 3, 4, or > 5 mismatches (indicated by different colours) with MiBird-U F/R primers. The total number of avian species evaluated was 410. The phylogenetic tree was constructed for species that can be amplified using MiBird-U primers (**b**). A total of 2,000 sequences were retrieved from the database to construct the phylogenetic tree. different classes are represented by filled circles with different colours. Lengths of branches correspond to the differences in sequences. Bar indicates edit distance.

The top BLAST hit with a sequence identity of at least 97% and E-value threshold of 10^-5^ was applied to species assignments of each representative sequence. Reliability of the species assignments was evaluated based on the ratio of total alignment length and number of mismatch bases between the query and reference sequences. For example, if a query sequence was aligned to the top BLAST hit sequence with an alignment length of 150 bp with one mismatch present, the ratio was calculated as 150/(1 + 1). The value one was added to the denominator to avoid zero-divisors. This value (e.g., 150/(1 + 1)) was calculated for the top and second-highest BLAST hit species, and the ratio score between these values was used as a comparable indicator of the species assignment. Results from the BLAST searches were automatically tabulated, with scientific names, common names, total number of reads and representative sequences noted in an HTML format. The above bioinformatics pipeline from data pre-processing through taxonomic assignment is available in supplements in a previous study1. Also, the above bioinformatic pipeline can be performed on a website. For more detailed information, please see http://mitofish.aori.u-tokyo.ac.jp/mifish. Please note that the pipeline implemented in the website currently uses the custom fish database and does not aim to detect avian species (confirmed on 20 September 2017).

## III. RESULTS AND DISCUSSION

### A. Tests of versatility of designed primers in silico and using extracted DNA

First, the performance of MiBird-U primers was tested *in silico* Fig.1s and Table 4). When G*/*T pairs were accepted, MiBird-U-F and -R perfectly matched 390 (95.1%) and 388 (94.6%) species among 410 species tested, respectively, and 99.5% and 96.8% of the 410 species showed at most 1 mismatch (Fig.1a). In addition, inter-specific differences in the edit distance were calculated and 82,177 out of 82,621 combinations (99.5%) showed edit distance larger than 5 (Table 4). To examine the range of species that can be amplified using MiBird-U primers, we performed an analysis with the primerTree package [25]. The results confirmed that the primers can amplify avian species (Fig.1b). MiBird-U primers can also amplify a diverse group of mammalian species in addition to amphibian, reptil ian and fish species (Fig.1b), which is not surprising because MiBird-U primers were produced by modifying fish/mammal-targeting universal primers. The capacity of MiBird-U primers to detect mammalian and other species might be useful when simultaneous detection of these animals is desired (e.g., when one tries to study co-occurrence patterns and potential interactions among animals).

**Table 4:** Frequency distributions of the interspecific edit distances of the primer set against bird sequences

| Edit distance | 0 | 1 | 2 | 3 | 4 | > 5 | Total |
| --- | --- | --- | --- | --- | --- | --- | --- |
| Inter-species | 27 | 50 | 67 | 113 | 187 | 82,177 | 82,621 combinations |
| Inter-genus | 10 | 23 | 40 | 80 | 157 |  |  |

Pairwise inter-species edit distances (= "Inter-species" row) were calculated for all species pairs, and pairwise inter-genus edit distances (= "Inter-genus" row) were calculated for pairs of species belonging to different genera

Second, the performance of MiBird-U primers was evaluated using 22 extracted avian DNA samples. All of the extracted DNA samples were successfully amplified, and the resultant sequences were deposited in the DDBJ*/*EMBLE*/*GenBank databases (Table 2). Together, the results of *in silico* tests and the amplification of extracted DNAs suggested that MiBird-U primers are capable of amplifying/identifying DNA fragments derived from diverse avian species.

### B. Primer testing with eDNA from field water samples

MiSeq sequencing and data pre-processing generated 656,675 sequences from 21 samples (including 3 field negative controls and 2 PCR negative controls) (Table 5). In general, the quality of sequences produced by our experiment was high (i.e., most raw reads passed the filtering process; Table S2).

**Table 5:** Sequence reads of detected species from water samples collected in the zoo

| Bird name living in cage<br>(scientific name) | The number of sequences of detected bird species |  |  |  |  |  |  |  |  |  |  |  |  |  |  |  | Non-target | Total |
| --- | --- | --- | --- | --- | --- | --- | --- | --- | --- | --- | --- | --- | --- | --- | --- | --- | --- | --- |
|  | *Haliaeetus* | *Larus* | *Tetrao* | *Chrysolophus* | *Tadorna* | *Tragopan* | *Goura* | *Aix* | *Spheniscus* | *Bubo* | *Ciconia* | *G. vipio* | *G. grus* | *Bucorvus* | *Parabuteo* | *Dromaius* | sequences* |  |
| Steller's sea eagle<br>(*Haliaeetus pelagicus*) | **28,448** | 0 | 0 | 0 | 0 | 0 | 0 | 0 | 0 | 0 | 0 | 0 | 0 | 0 | 0 | 0 | 3,238 | 31,686 |
| Black-tailed gull<br>(*Larus crassirostris*) | 0 | **4,437** | 0 | 0 | 0 | 0 | 0 | 0 | 0 | 0 | 0 | 0 | 0 | 0 | 0 | 0 | 17,922 | 22,359 |
| Capercaillie<br>(*Tetrao urogallus*) | 0 | 0 | **36,095** | 0 | 0 | 0 | 0 | 0 | 0 | 0 | 0 | 13 | 0 | 0 | 0 | 0 | 19,477 | 55,585 |
| Lady Amherst's pheasant<br>(*Chrysolophus amherstiae*) | 0 | 0 | 0 | **39,151** | 0 | 0 | 0 | 25 | 0 | 0 | 0 | 0 | 15 | 0 | 0 | 0 | 8,130 | 47,321 |
| Ruddy shelduck<br>(*Tadorna ferruginea*) | 0 | 0 | 0 | 138 | **2,848** | 209 | 2,750 | 7,939 | 0 | 0 | 0 | 0 | 0 | 0 | 0 | 0 | 12,498 | 26,382 |
| Temminck's tragopan<br>(*Tragopan temminckii*) | 0 | 0 | 376 | 0 | 0 | **57,072** | 0 | 0 | 0 | 0 | 0 | 0 | 0 | 0 | 0 | 0 | 1,944 | 59,392 |
| Victoria crowned pigeon<br>(*Goura victoria*) | 11,023 | 0 | 0 | 0 | 0 | 0 | **2,186** | 24 | 0 | 0 | 0 | 0 | 0 | 0 | 0 | 0 | 13,108 | 26,341 |
| Mandarin duck<br>(*Aix galericulata*) | 254 | 0 | 0 | 0 | 65 | 51 | 34 | **13,465** | 0 | 0 | 0 | 0 | 0 | 0 | 0 | 0 | 12,272 | 26,141 |
| Humboldt penguin<br>(*Spheniscus humboldti*) | 1,905 | 0 | 0 | 0 | 289 | 0 | 85 | 428 | **1,834** | 0 | 0 | 0 | 0 | 0 | 0 | 0 | 28,091 | 32,632 |
| Snowy owl<br>(*Bubo scandiacus*) | 901 | 0 | 0 | 0 | 65 | 0 | 64 | 191 | 0 | **425** | 0 | 0 | 0 | 0 | 0 | 0 | 28,566 | 30,212 |
| Oriental white stork<br>(*Ciconia boyciana*) | 7,258 | 63 | 0 | 0 | 0 | 0 | 103 | 258 | 0 | 0 | **3,072** | 0 | 0 | 0 | 0 | 0 | 22,815 | 33,569 |
| White-naped crane<br>(*Grus vipio*) | 186 | 0 | 527 | 0 | 22 | 0 | 36 | 33 | 0 | 0 | 0 | **59,678** | 0 | 31 | 0 | 0 | 2,390 | 62,903 |
| Common crane<br>(*Grus grus*) | 0 | 0 | 0 | 548 | 0 | 0 | 0 | 0 | 0 | 0 | 0 | 0 | **52,717** | 0 | 0 | 0 | 2,586 | 55,851 |
| Southern ground hornbill<br>(*Bucorvus cafer*) | 148 | 0 | 0 | 0 | 0 | 15 | 0 | 0 | 0 | 0 | 0 | 0 | 0 | **36,955** | 0 | 0 | 4,900 | 42,018 |
| Harris's hawk<br>(*Parabuteo unicinctus*) | 1,227 | 0 | 0 | 0 | 77 | 0 | 0 | 332 | 0 | 0 | 0 | 0 | 0 | 0 | **306** | 0 | 20,338 | 22,280 |
| Emu<br>(*Dromaius novaehollandiae*) | 396 | 0 | 0 | 0 | 67 | 0 | 25 | 146 | 0 | 0 | 0 | 22 | 0 | 0 | 0 | **1,647** | 3,412 | 5,715 |
| Field NC | 0 | 0 | 0 | 0 | 0 | 0 | 0 | 0 | 0 | 0 | 0 | 0 | 0 | 0 | 0 | 0 | 27,218 | 27,218 |
| Field NC | 0 | 0 | 0 | 0 | 0 | 0 | 0 | 0 | 0 | 0 | 0 | 0 | 0 | 0 | 0 | 0 | 7,977 | 7,977 |
| Field NC | 0 | 0 | 0 | 0 | 0 | 0 | 0 | 0 | 0 | 0 | 0 | 0 | 0 | 0 | 0 | 0 | 40,890 | 40,890 |
| PCR NC | 0 | 0 | 0 | 0 | 0 | 0 | 0 | 0 | 0 | 0 | 0 | 0 | 0 | 0 | 0 | 0 | 0 | 0 |
| PCR NC | 0 | 0 | 0 | 0 | 0 | 0 | 0 | 0 | 0 | 0 | 0 | 0 | 0 | 0 | 0 | 0 | 0 | 0 |
| Total sequence | 51,746 | 4,500 | 36,998 | 39,837 | 3,433 | 57,347 | 5,283 | 22,841 | 1,834 | 425 | 3,072 | 59,713 | 52,732 | 36,986 | 306 | 1,647 | 277,772 | 656,472 |

\*See Table S3 for the content of non-target sequences. Bold numbers indicate the number of sequences of a target bird species.

Among the 16 water samples from zoo cages examined here, all avian species were successfully detected (Table 5). Briefly, eDNA samples of the Steller’s sea eagle (*Haliaeetus pelagicus*), capercaillie (*Tetraourogalluis*), white-naped crane (*Grus vipio*), common crane (*Grus grus*) and southern ground hornbill (*Bucorvus cafer*) generated high numbers of sequence reads, and 64.9−94.9% of total sequence reads were assigned to the target avian species. Samples from cages of the black-tailed gull (*Larus crassirostris*), Humboldt penguin (*Spheniscus humboldti*), snowy owl (*Bubo scandiacus*), oriental white stork (*Ciconia boyciana*), Harris’s hawk (*Parabuteo unicinctus*) and emu (*Dromaius novaehollandiae*) generated fewer sequence reads, and 1.4−28.8% of total sequence reads were assigned to the target avian species. The reason for these variations in the proportions of sequence reads from target avian species is not known, but as discussed in the previous study [9], the observed levels of variations were not surprising because detection of animals’ sequences relies on contacts of animals with water and because opportunities for animals to contact water would depend on animals’ behaviour. These considerations imply that the proportion of sequence reads from a particular avian species would be inherently spatially and temporally stochastic to some extent (see also results of mammalian eDNA metabarcoding in Ushio et al. [9]). It is not surprising that sequences of the Lady Amherst’s pheasant, ruddy shelduck, Temminck’s tragopan, Victoria crowned pigeon and mandarin duck were detected in the ruddy shelduck sample (Table 5) because all of these five species were kept in the bird cage where the ruddy shelduck sample was collected.

In addition to the target avian species, we frequently detected many non-target species (Table 5 and S3). For example, sequences of the Steller’s sea eagle were frequently detected in other samples, e.g., the Victoria crowned pigeon, oriental white stork, Humboldt penguin and so on (Table 5). As our field negative controls generated no target bird sequences (Table 5), it does not seem likely that the detection of the sea eagle in other samples was due to cross-contamination during sampling or experiments. One possible reason for the detection of non-target avian species include the spatial closeness of the eagle’s cage and the other cages. For instance, the cages of the Victoria crowned pigeon (i.e., the bird cage) and Humboldt penguin were located close to the eagle’s cage, and thus it is possible that the eagle’s feathers and other tissues could be transported (e.g., via wind) to other cages. Also, zoo staff frequently moved among cages, and they were possible transporters (e.g., through their shoe sole) of materials containing DNA of non-target species.

Other frequently detected non-target species were falcated teal (*Anas falcata*), common shelduck (*Tadorna tadorna*), common moorhen (*Gallinula chloropus*), fishes and humans (Table S3). The falcated teal, shelduck and moorhen were not kept in cages, but wild common moorhens and close relatives of the duck and shelducks (i.e., Eurasian wigeon [*Anas penelope*] and common pochard [*Aythya ferina*], respectively) are commonly observed in the sampling region (e.g., regulating ponds on-site), and thus their DNA might have contaminated zoo cages (possibly via feathers, feces or other tissues) and thus have been detected by the metabarcoding.

The frequently detected fish species here are also species that are commonly observed in Japan, and the zoo uses waters from a natural lake and rivers. Therefore, the fish sequences might have been derived from water under natural conditions. Detection of many human sequences was not surprising considering that visitors to the zoo and staff members, who are potential sources of human sequences, are almost always near the cages. It is also be pos-sible that contaminations of human and fish DNA happened under the laboratory conditions (Table S3), because in our lab fish DNAs were routinely processed and humans were often working. The sequences of these obvious non-target taxa (i.e., humans and fish) may be excluded from further statistical analyses if one may be interested in ecological interpretations of the results.

Lastly, in order to test the usefulness of MiBird primers under a natural field condition, we performed a metabarcoding study using a water sample from a pond adjacent to the Natural History Museum and Institute, Chiba. As a result of MiSeq sequencing, 14,873 reads of avian species were generated from three water samples, and five avian species (common shoveler [*Anas clypeata*], 883 reads; falcated teal, 3,246 reads; common moorhen, 9,260 reads; light-vented bulbul [*Pycnonotus sinensis*], 745 reads; and common shelduck, 739 reads) were detected. As a systematic monitoring of the bird community (e.g., frequent visual observation) has not been performed in the study site, rigorous validation of the metabarcoding study was not possible. Some avian species detected, i.e., light-vented bulbuls, common shelducks and falcated teals, are rare, or not reported, in this region, suggesting that these species were misidentified. These possible misidentifications are likely to be attributable to a lack of reference sequences and/or insufficient inter-species differences in the amplified DNA region (i.e., partial 12S mitochondrial region). Light-vented bulbuls, common shelducks and falcated teals are relatives of brown-eared bulbuls (*Hypsipetes amaurotis*), common pochards (*Aythya ferina*) and Eurasian wigeons (*Anas penelope*), respectively, and these relatives are indeed common inhabitants in the sampling region. Together, these results suggest that MiBird primers were capable of detecting bird species under a field condition, but at the same time, improvements of reference sequence databases, further validations of MiBird primers, and careful interpretations are necessary.

## IV. CONCLUSION

The present study demonstrated that eDNA analysis combined with our new primer sets and the MiSeq platform can be a useful tool for detecting bird species. Describing and monitoring the diversity of bird species, as well as other animals, is one of the critical steps in ecosystem conservation and management, but it can be laborious, costly and incomplete if one relies on a few traditional survey methods. The eDNA metabarcoding approach presented here is non-invasive and efficient. Moreover, as information of non-target organisms (e.g., invertebrates and microbes in our case) is also encoded in eDNA, analyzing eDNA of organisms from multiple taxa might be useful for studying co-occurrence patterns and even potential interactions among organisms (e.g., bird-insect interactions). In conclusion, we propose that the eDNA metabarcoding approach can serve as an efficient alternative for taking a snapshot of bird diversity and could potentially contribute to effective ecosystem conservation and management.

### Ethics

This study was approved by Yokohama Zoological Gardens ZOORASIA.

### Authors’ contributions

MU and MM conceived and designed research; MU, IN and KM performed sampling; MU, IN, TS and MM performed experiments; MU, MT and WI performed data analysis; MU and MM wrote the early draft and completed it with significant inputs from all authors.

## Acknowledgements

We would like to thank Noriya Saito, assistant manager of Yokohama Zoological Gardens ZOORASIA for help in sampling at the zoo, and Asako Kawai for assistance with experiments. We also thank Hiroki Yamanaka of The Department of Environmental Solution Technology/The Research Center for Satoyama Studies in Ryukoku University for providing the opportunity for us to use the Illumina MiSeq platform. This research was supported by PRESTO (JPMJPR16O2) from Japan Science and Technology Agency (JST), CREST (JPMJCR13A2) from Japan Science and Technology Agency (JST), and ERTDF (4-1602) The Environment Research and Technology Development Fund, Japan.

